# Targeting Histamine H₃ Receptors to Suppress NGF-Driven Hyperexcitability in Neuropathic Pain

**DOI:** 10.64898/2026.08.28.746833

**Authors:** Ajeet Kumar, Rakesh Kumar, Poonam Kumari, Aditi Bagade, Anjali Yadav, Boda Arun Kumar, Shalini Dogra, Deepmala Umrao, Bryan Copits, Charles F. Zorumski, Steven J. Mennerick, Prem N Yadav

## Abstract

Peripheral nerve injury induces long-lasting changes in sensory neurons that contribute to the development of neuropathic pain. Although the histamine H₃ receptor (H₃R) is best known for regulating neurotransmitter release in the central nervous system, its expression and role in dorsal root ganglia (DRG) remain poorly understood. In this study, we examined the expression, cellular distribution, and functional role of H₃R in DRG and implications in neuropathic pain. Sciatic nerve injury increased membrane-associated H₃R protein expression in the DRG, suggesting enhanced receptor trafficking or stabilization at the neuronal membrane. Single-cell transcriptomic analysis revealed that H₃R mRNA expression in DRG is predominantly restricted to peptidergic sensory neurons and C-LTMRs rodents. Fluorescence imaging and transcriptomic analysis also suggest that the majority of H3R co-express with TrkA^+^ (a NGF receptor) sensory neuron populations. These findings identify a subset of NGF-responsive nociceptors in which H₃R may directly influence injury-induced sensitization. Pain behavior assessment demonstrated a paradoxical role of an H3R inverse agonist (GSK334429), which reduces mechanical hypersensitivity caused by chronic constriction injury (CCI) with no change in thermal pain. Patch-clamp recordings show that the GSK334429 attenuates NGF-induced hyperexcitability in DRG neuronal cultures. Overall, our work suggests a contributing role of H3R in modality-specific effects on sensory processing through its enrichment in specific neuronal populations in DRG.

**Graphical Abstract:** 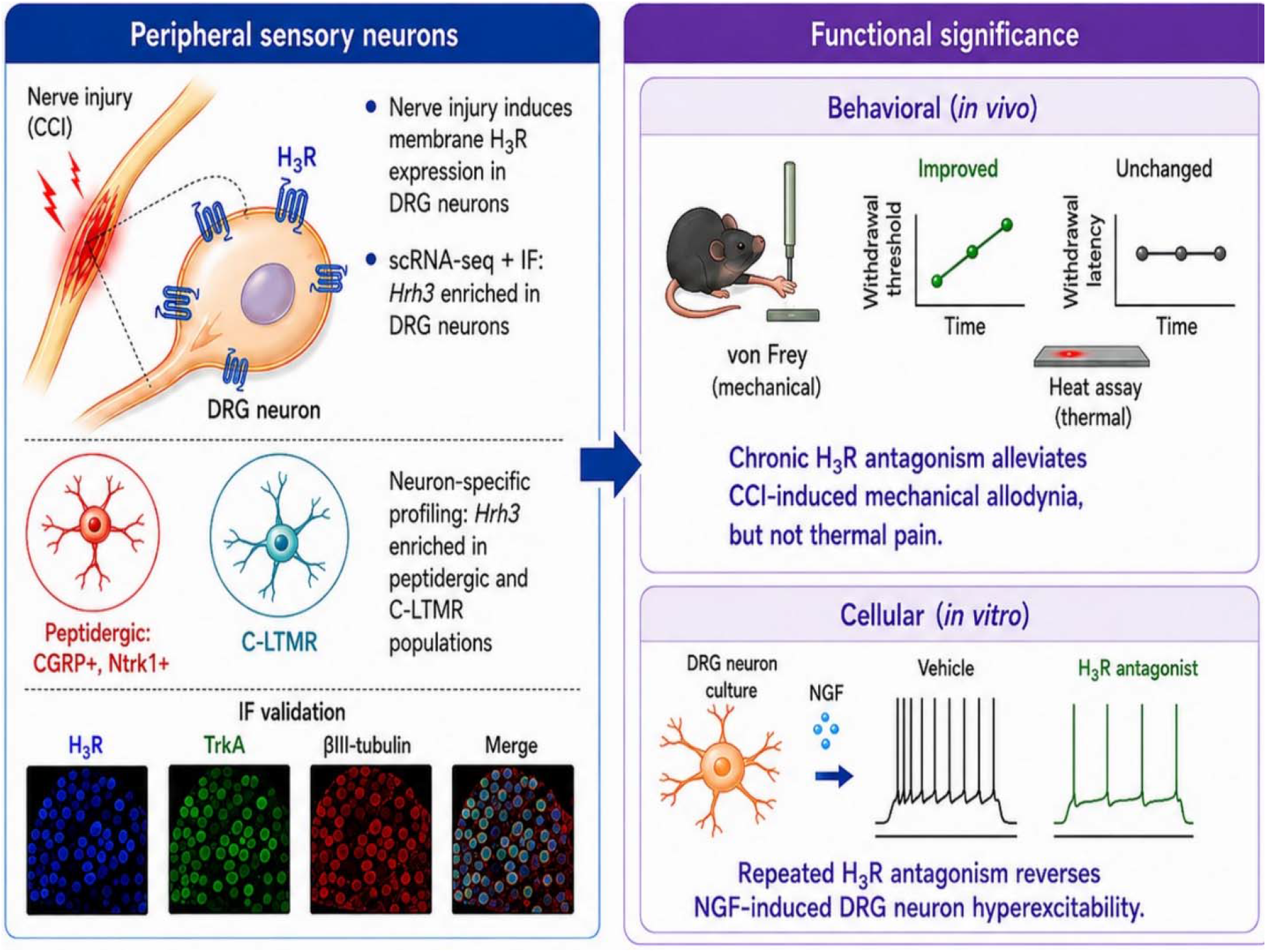

## 1. Introduction

The dorsal root ganglion (DRG) comprises subpopulations of sensory neurons and supporting cells that play roles in detecting and transmitting somatosensory and nociceptive signals. Various single-cell transcriptomic studies have expanded our understanding of distinct somatosensory neuron classes, including peptidergic and non-peptidergic nociceptors, myelinated mechanoreceptors, proprioceptors, and specialized populations such as C low-threshold mechanoreceptors (C-LTMRs), which have characteristic structures and functions (Usoskin et al., 2015; Li et al., 2016). These primary afferent fibers relay sensory information to specific regions of the dorsal horn, where second-order neurons then project this to different areas of the brain. However, nerve injury lowers threshold for DRG neuron firing, through multiple mechanisms involving inflammation and alterations in gene expression pathways, resulting in persistent hypersensitivity to non-noxious signals (Gold and Gebhart, 2010; Djouhri et al., 2012; Jang and Garraway, 2024). Among the DRG primary sensory neurons, peptidergic nociceptors are of interest because they are injury responsive and express multiple genes that are associated with injury induced hypersensitization. These include tropomyosin receptor kinase A (TrkA) (encoded by NTRK1), which, upon binding nerve growth factor (NGF), triggers profound hypersensitivity (Hirose et al., 2016; Hu et al., 2016; Norman and McDermott, 2017).

Neuropeptides released during heightened inflammation following nerve injury serve as mediators in driving sustained hypersensitivity. For example, NGF participates in retrograde pain signaling to the DRG by binding to TrkA and activating downstream signaling cascades, including PI3K–AKT, MAPK/ERK, and Src–STAT3 (Delcroix et al., 2003; Barker et al., 2020). This results in activation of voltage-gated sodium channels and upregulation of CGRP, thereby enhancing excitability and peripheral sensitization (Di Castro et al., 2006; Schaefer et al., 2018). Alongside NGF, histamine is an injury-associated inflammatory mediator (Woolf, 2011; Rosa and Fantozzi, 2013; Akiyama et al., 2014; Sommer et al., 2018) that regulates DRG neurons activity and spinal dorsal horn signaling through cognate histamine G-protein coupled receptors (Cannon et al., 2007; Akiyama et al., 2014). Thus, in addition to NGF-responsive pain signaling, histamine signaling is an important target for modulating peripheral sensitization after nerve injury.

Histamine mediates inflammation and neurotransmission by binding to four receptor isoforms: H₁R, H₂R, H₃R, and H₄R (Dong et al., 2014; Panula et al., 2015). Amongst them, Gα_i_-coupled H₃R functions both as an auto-receptor by inhibiting the synthesis and release of histamine from histaminergic neurons and as a heteroreceptor by suppressing the release of other neurotransmitters from non-histaminergic neurons (Panula et al., 2015; Nieto-Alamilla et al., 2016). Importantly, activation of H_3_R inhibits voltage-gated sodium channels at synapses, thereby suppressing calcium influx and preventing neurotransmitter release (Brown et al., 2001; Vázquez-Vázquez et al., 2022). The actions of H_3_R activation are unusual among typical pro-inflammatory effects of histamine, in that they may work against pro-excitatory effects associated with hypersensitivity. Additionally, H3R signaling, particularly PLCβ mediated PIP2 hydrolysis may converge with NGF/TrkA signaling and promote nociceptor excitability by suppressing potassium channel activation (Chuang et al., 2001; Linley et al., 2008).

Despite a plethora of studies on H_3_R in the central nervous system (Hough and Rice, 2011), little is known about its distribution and its role within DRG remains poorly understood. Although several studies have demonstrated that both H_3_R antagonists and inverse agonists reduce neuropathic pain in rodent models, the underlying mechanisms remain unclear in light of the physiological actions (Medhurst et al., 2008; McGaraughty et al., 2012; Łażewska and Kieć-Kononowicz, 2018; Popiolek-Barczyk et al., 2018; Degutis et al., 2025). Here we hypothesize that H_3_Rs may represent a therapeutically relevant target on subpopulations of DRG neurons through pro-excitatory H3R pathways. To address these questions, we quantified H_3_R membrane protein in DRG with and without nerve injury. We also reanalyze publicly available single-cell RNA sequencing datasets and found Hrh3 (H_3_R) enrichment in peptidergic nociceptors and C-LTMR sensory neurons. To determine whether H3R upregulation in a neuropathic pain model is protective or promotes pain sensitivity, we examined the effects of the H3R inverse agonist GSK334429 (GSK). We evaluated neuropathic pain behaviors in vivo and performed whole-cell patch-clamp recordings from cultured DRG neurons to assess whether the compound alters NGF-induced neuronal hypersensitivity.

Our findings demonstrate a paradoxical role of H_3_R; GSK reduced mechanical hypersensitivity caused by chronic constriction injury (CCI), with no change in thermal sensitivity and expression of thermal transduction mediator TRPV1. Patch-clamp recordings show that the inverse agonist also normalized NGF-induced hyperexcitability of DRG neurons in culture. Overall, our work suggests contributing role of H_3_R in modality-specific effects on sensory processing, through its enrichment in specific neuronal populations in DRG.

## 2. Methods and materials

### 2.1 Animals

All animal experiments including behavior, CCI surgery procedures were conducted in accordance with the Guide for the Care and Use of Laboratory Animals and were approved by the Institutional Animal Ethics Committee (IAEC) of CSIR-Central Drug Research Institute, Lucknow, India. A total of 40 male Sprague Dawley (SD) Rats (6-8 weeks old and with 200-250g body weight) were used in this study. Animals were housed (4 rats/cage) on a 12-hour (h) light/dark cycle with the lights turned on at 8.00 am. Additionally, strict control was placed on the room environment (22±2 °C) and humidity (50-70%). Food pellets (Altromin International, Germany, Cat. No. 1320) and water (autoclaved sterile) were provided ad *libitum*. Primary DRG cells culture procedures were conducted in accordance with the National Institutes of Health guidelines and approved by the Washington University Institutional Animal Care and Use Committee.

### 2.2 Chronic constriction injury of the sciatic nerve

To study chronic neuropathic pain, we established a unilateral sciatic nerve CCI model for *in-vivo* experiments (Austin et al., 2012). In brief, the animals underwent anesthesia by intraperitoneal injections (*i.p.*) of ketamine and xylazine: 90mg/kg &10mg/kg, respectively. The sciatic nerve was exposed following a skin incision by separating the connective tissues between the gluteus superficialis and biceps femoris muscles. At 1 mm intervals, four loose ligatures (silk sutures, 5.0) on the sciatic nerve were made around the sciatic nerve to induce neuropathy. The connective tissue was brought back close, and the skin incision was sutured with silk sutures (3.0). The animals were allowed to regain consciousness and then transferred back to the animal facility.

### 2.3 Study design and treatment regimen

To evaluate pain behavior with H_3_R antagonist in neuropathic pain model we used GSK334429 hydrochloride (Sigma-Aldrich; Cat #SML0500). GSK solution was prepared by dissolving the powder in normal saline. The solution was administered at a dose of 0.5 mg/kg body weight to rats via *intraperitoneal* (*i.p.*) injection. Following CCI surgery on Day 1, mice received GSK twice daily from Day 5 through Day 13.

### 2.4 Behavioral procedure

The study used 40 SD Rats in two separate CCI-induced neuropathic pain experiments. To remove biases, all behavioral experiments were conducted at the same time of the day.

#### 2.4.1 von Frey Hair test

We followed the well-established test to assess mechanical sensitivity or response as described by Maximilian von Frey (Pitcher et al., 1999). In brief, classical von Frey filaments (made of nylon monofilaments with different bending forces; Stoelting, Cat#58011) were manually applied on the midplantar region of mice’s hind paw in ascending order of stiffness. For the same reason, the mice were caged in a transparent box with a metallic mesh base. The filaments were applied a total of 5 times until brisk withdrawal of the paw, indicating pain induction in both control and experimental mice. The threshold value (grams) was defined as the filament resulting in left hind paw withdrawal three times out of 5 trials. Baseline mechanical sensitivity was assessed on Day 0 before CCI surgery. Following CCI, mechanical sensitivity was evaluated every second day immediately before the morning dose (10–12) hours after the previous injection) to assess the sustained analgesic effect, and acute effect evaluated after 30 min after drug. Sham control animals received vehicles according to the same treatment and testing schedule.

#### 2.4.2 Hot plate test

Thermal nociception is a well-established assay for evaluating supraspinal pain responses to heat stimuli. We assessed thermal pain using the hot plate test. Within an open-topped, transparent rectangular box (30cm high), a 12 × 27 cm aluminum plate was maintained at a constant temperature of 50°C. Plate temperature was regulated via electronic proportional feedback circuit and start/stop/reset functions were controlled using a remote foot-switch pad. The animals were acclimatized to the behavioral room and temperature conditions (22 ± 2°C) by transferring them a day before the experimental day. The tests were conducted at the same time, 09:00 and 17:00 h, to avoid variability. The rats were gently placed on the preheated plate for each trial, and the latency to response was recorded as the first hind-paw lick or jump. We set a 120-second cut-off time to prevent injuries to the animals.

### 2.5 Wheat germ agglutinin (WGA) membrane protein pull-down

To assess H_3_R expression in DRG tissues, WGA pull-down assay was performed as previously described (Sona et al., 2018). In brief, DRG were pooled to prepare a lysate, and 150 μg of protein lysate from each condition was incubated with 40 μl of WGA beads at 4°C for 6 hours with rotation in the cold room to allow sufficient binding of glycosylated membrane proteins to the WGA beads. The beads were washed three times with wash buffer for 10 min each (1X phosphate-buffered saline containing 0.1% Triton X-100 and 1X protease inhibitor cocktail). Later, elution of the bound proteins was performed by using 30 μL of 2X Laemmli SDS-PAGE sample buffer (0.1% 2-mercaptoethanol, 0.0005% bromophenol blue, 10% glycerol, 2% SDS, and 50 mM Tris-HCl, pH 6.8). Equal amounts of both WGA-enriched membrane protein and respective input lysate samples (30 μg) were loaded onto SDS-PAGE and immunoblotted with an anti-H_3_R antibody (1:2000, RRID: AB_1587114; Millipore) to assess H_3_R protein expression. In addition, the membrane was probed with β-actin (1:10000; RRID: AB_476697; Sigma Aldrich) to ensure equal sample loading and for further normalization during quantification.

### 2.6 Western blotting

Protein lysates were prepared from DRG tissues as previously described (Dogra et al., 2016a), and both lysates were utilized for performing a western blot. Briefly, the sample was prepared by using 50 μg protein in 4X Laemmli buffer, followed by denaturation at 95° C heat block. After a short spin of the samples, the proteins were loaded onto 8-12% polyacrylamide gels and resolved by SDS-PAGE, initially at 80V and then at 120V. After the proteins resolved, they were transferred to polyvinylidene difluoride (PVDF) membranes (pre-wetted with methanol) in transfer buffer (Tris-glycine-methanol) at 100 V for 1 hr. Membranes were blocked for 2 hours at room temperature in blocking buffer containing 10% bovine serum albumin (BSA) in 1X Tris-buffered saline (TBS), followed by overnight incubation with primary antibodies (prepared in TBST) at 4°C. Next day, the membranes were washed three times with TBST for 10 mins each, followed by 2 h incubation with horseradish peroxidase (HRP)-conjugated secondary antibodies at room temperature. After three additional washes, the membranes were probed with enhanced chemiluminescence (ECL) solution (Merck-Millipore, India) to detect protein bands. The bands were visualized using a gel documentation system (MyECL Imager, Thermo Fisher Scientific, USA). The analysis was performed by densitometric quantification of band intensities using ImageJ software.

### 2.7 Immunofluorescence (IF) staining

Lumbar DRG tissue sections were collected from 6-8 weeks old SD rats to perform immunohistochemistry, or the lumbar DRG neurons were cultured to perform immunocytochemistry as previously described (Dogra et al., 2016a). The tissues were fixed in 4% paraformaldehyde (PFA) overnight, then transferred to 30% sucrose solution in phosphate buffer. Tissues were embedded in OCT medium in a cryomold and stored at −20 °C until use. The tissue was cryosectioned into 10 μm sections. The slides were stored at −20 °C. Slides to be used for immunostaining were equilibrated at RT for 15 mins before washing with PBS. Each of the three PBS washes was carried out for 10 minutes, with slides kept in the slide chamber on the shaker.

*For in vitro studies*, DRG neurons were cultured on poly-L-lysine-coated coverslips. On days 6 or 8, the media was discarded from the neuron culture, and the cultures were rinsed prior to fixation with 4% PFA for 10 minutes at 4 °C. The neurons were washed with PBS three times, with each wash for 5 minutes. After PBS washes, the steps remained the same for both tissue sections and the cultured DRG cells. Blocking was carried out with 5% normal goat serum in 1X PBST (1× PBS + Triton X-100: 0.3% for tissues, 0.1% for cultures) for 1.5 hours at room temperature, followed by incubation with primary antibodies prepared in 5% blocking solution (anti-H_3_R, 1:500, RRID:AB_10001669; TrkA, 1:500, RRID:AB_2283049; NeuN, 1:2000, RRID:AB_2298772; GFAP, 1:2000, RRID:AB_11212597; β-Tub3, 1:1500, RRID:AB_477590; NF-200, 1;1000, RRID:AB_477257; CGRP, 1:500, RRID:AB_2290729 and IB4, 1:500, RRID:SCR_014365) overnight at 4 °C. The next day, the samples were washed with PBST to remove any residual primary antibody. Later, depending on the host of primary antibodies, the appropriate fluorophore-conjugated secondary antibodies were used for 1 hour at room temperature. The samples were washed three times with PBST, with each wash for 5 minutes to remove any residual traces of secondary antibodies. The slides and coverslips were mounted with medium containing DAPI (Cat. No. H-1000, Vector Laboratories). The images were captured using a confocal microscope at 20X magnification.

### 2.8 Transcriptomic profiling

Single-cell RNA-seq datasets were retrieved from the NCBI Gene Expression Omnibus (GEO), including GSE174430 of adult mouse DRG (Jager et al., 2022), GSE139088 from embryonic E11.5 DRG (Sharma et al., 2020), and GSE254789 from transcriptionally defined adult sensory neuron subtypes (Qi et al., 2024). These datasets were analyzed to investigate gene expression patterns across developing and mature sensory neuron populations. Because these datasets were provided in different file formats, including 10x Genomics Matrix Market files, H5-based expression matrices, count matrices, and tab-delimited count/metadata tables, each dataset was imported into Python. Files were then converted into compatible AnnData object for downstream analysis. We retained the information on the sample, condition, and cell population as specified in the metadata. As required for the combined analysis of DRG, datasets were integrated; otherwise, they were used in isolation.

All datasets were processed entirely in Python using the Scanpy workflow (Wolf et al., 2018). For each cell, we used total transcript counts, the total number of detected genes, and the percentage of mitochondrial genes as quality-control parameters. Threshold filtering allowed us to remove low-quality cells within each dataset. To account for data variance and differences in sequencing depth across cells, standard preprocessing was performed: library-size normalization to 10,000 counts per cell and logarithmic transformation. This allowed us to extract information on highly variable genes, regress out unwanted variation due to total gene or transcript counts, mitochondrial percentage, and data scaling, and to scale the data. We used principal component analysis to filter out the noise from actual biologically significant signals. Before clustering cells into distinct cell types using Leiden clustering, a nearest-neighbor graph was constructed, and Uniform Manifold Approximation and Projection (UMAP) was used to project biologically significant information into 2-dimension. Cell-specific gene markers were used to assign cell-type identity to neurons, satellite glia, Schwann cells, endothelial cells, fibroblasts, and immune cells of DRG. Similarly, subtype markers were used to further evaluate DRG neuronal subclasses.

Finally, we determined the H_3_R expression pattern in DRG sensory neuronal populations. Additionally, we also evaluated the expression pattern of histamine- and other neurotrophin-related genes across DRG cell types to determine if there is an association between H_3_R and histaminergic or neurotrophins signaling pathways. To study histaminergic signaling pathways, we analyzed the expression of genes including *Hdc*, *Hrh1* (H_1_R), *Hrh2* H_2_R), *Hrh3* (H_3_R), *Hrh4* (H_4_R), *Slc22a3*, *Slc18a2*, and *Maob*, which are associated with histamine synthesis, receptors, metabolism, and transport. Similarly, to study neurotrophin signaling pathways, we assessed the expression of genes including Ngf, *Bdnf*, *Ntf3*, *Ntrk1*, *Ntrk2*, *Ntrk3*, and *Ngfr*. We first confirmed their availability and then examined the expression patterns of these genes using a dot or UMAP feature plot in DRG datasets. We also assessed the co-expression of H_3_R and TrkA across DRG cell populations including neuronal subtypes.

### 2.9 Primary DRG neuronal culture

DRG cultures were prepared from 1- to 2-day-old wild-type C57BL/6J mice as described previously (Copits et al., 2021). Mice were anesthetized with isoflurane, spinal columns were removed, and DRG were dissected in HBSS + 10 mM HEPES (HBSS+H). Tissue was enzymatically digested with papain (45U, Worthington, cat. # LS003126) for 20 min at 37°C, rinsed in HBSS+H, followed by collagenase (1.5 mg/mL; Sigma, cat. #C6885) for 20 min at 37°C. Ganglia were washed and resuspended in culture media consisting of Neurobasal A (GIBCO), 5% FBS (Life Technologies), 1x B27 (GIBCO), 2 mM glutamax (Life Technologies), and 100 μg/mL penicillin/streptomycin (Life Technologies). Neurons were dissociated by mechanical trituration through glass pipettes and filtered through 40 μm filters. Isolated DRG neurons were resuspended and seeded at a density of 5,000 cells/well onto 12-mm glass coverslips coated with poly-D-lysine and collagen. DRG neurons were maintained in culture in Dulbecco’s MEM/F12 (1:1) supplemented with 10% FBS (Thermo Scientific) and penicillin (100 IU/ml) and streptomycin (100 μg/ml; Life Technologies) for 5 to 6 days before experiment.

### 2.10 Patch-clamp electrophysiology

DRG neurons were treated at DIV 5/6 with NGF 100 ng/mL (Alomone Labs Cat# N-100) in the presence or absence of GSK (10 µM; single or double treatment), with vehicle-treated cultures serving as controls. After 48 h of treatment, DRG neurons at DIV 7/8 were transferred to the recording chamber mounted on an Eclipse TE2000S inverted microscope. Whole-cell patch-clamp recordings were performed using a Multiclamp 700B amplifier, digitized with an Axon Digidata 1550 Low-Noise Acquisition System, and acquired and analyzed using pClamp 10 software (Molecular Devices). For action potential recordings, the intracellular pipette solution contained the following, in mM: 120 K-gluconate, 5 NaCl, 3 MgCl₂, 10 HEPES, 1.1 EGTA, 0.1 CaCl₂, 4 Na₂ATP, and 0.4 Na₂GTP; the pH was adjusted to 7.25 with KOH. The extracellular recording solution typically contained the following, in mM: 145 NaCl, 3 KCl, 10 HEPES, 10 glucose, 2 CaCl₂, and 1 MgCl₂, pH 7.25 and Osmolality 300-310mOsm/kg. Patch pipettes were pulled from borosilicate glass capillaries and had resistances of 3–6 MΩ. Membrane properties and neuronal excitability were assessed in current-clamp mode. After establishing a stable whole-cell configuration, resting membrane potential and spontaneous activity were recorded for 1-min under gap-free current-clamp conditions without current injection. Membrane excitability was then evaluated using 1-s stepwise current injections ranging from −50 pA to +140 pA (Δ10 pA). Rheobase was defined as the minimum current injection required to evoke an action potential. Spike firing was quantified as the number of action potentials generated in response to each current step.

## 3. Results

### 3.1 Nerve injury increases membrane-associated H₃R expression in DRG

Multiple previous studies have shown that H₃Rs can be targeted to alleviate neuropathic pain by antagonists and inverse agonists (Chaumette et al., 2018; Łażewska and Kieć-Kononowicz, 2018; Obara et al., 2020); however, whether chronic peripheral nerve injury alters H₃R expression in DRG sensory neurons and glial cells remains unknown. To assess H₃R protein levels in DRG, we used CCI of the sciatic nerve as a model of neuropathic pain in rats. First, the CCI model was validated by assessing mechanical pain before and after surgery. Paw withdrawal threshold was measured using von Frey hair test. As expected, CCI of the sciatic nerve significantly reduced paw withdrawal threshold in the ipsilateral side compared to the sham-operated animal at post-surgery time points (5, 9, and 14 days), (**Figure 1A**, Two-way ANOVA *p < 0.05, **p < 0.01, N=6/group). Two weeks after surgery, we collected L4-L5 DRGs from the ipsilateral side of CCI and sham-operated rats. Glycosylated membrane proteins from DRG lysate were obtained by WGA pull-down assay to examine the expression levels of membrane-associated H₃R by western blot analysis. To ensure equal input, β-actin was used as a loading control from the flow-through fraction (non-glycosylated cytosolic proteins) (Figure 1B). Western blotting from the WGA-enriched fractions with H₃R antibody revealed CCI increases H₃R levels in DRG tissues compared to sham controls (**Figure 1B**), and quantitative analysis normalized with respective β-actin indicated that CCI significantly upregulated H_3_R expression compared to sham controls (**Figure 1C**, unpaired Student’s t-test; **p < 0.01 N=5). To further validate that the pulled-down proteins are enriched in membrane proteins, we assessed the levels of the transferrin receptor (TfR), a transmembrane glycoprotein, on the same WGA-enriched blot. The expression levels of TfR did not differ between the CCI and sham groups, supporting the quality and consistency of the membrane protein isolation procedure (**Figure 1 C).**

**Figure 1:**
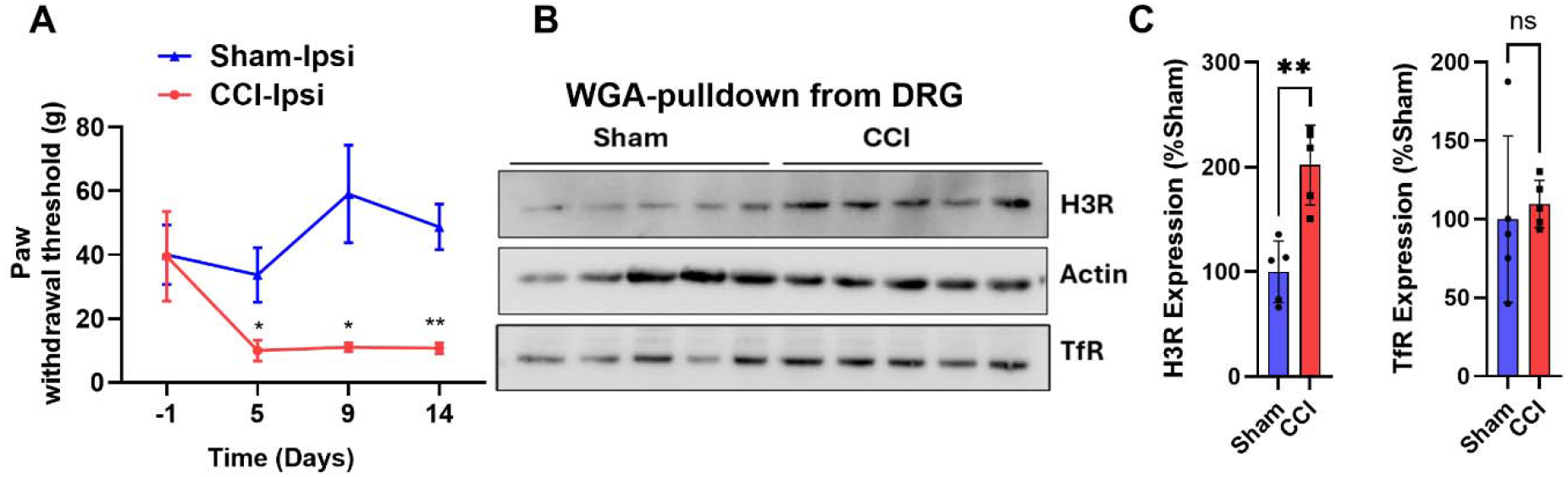
Histamine 3 receptor expression increases in DRG following chronic constriction injury. (A) Animals that underwent CCI showed robust mechanical hypersensitivity compared to sham controls as measured by mechanical nociceptive thresholds using the von Frey filament assay (Two-way ANOVA F _(1,10)_ =12.90, *p < 0.05, **p < 0.01, N = 6) (B) Representative Western blots for H_3_R expression in WGA-pulldown DRG lysates from sham and CCI animals. β-actin represents a loading control, and TfR as a control for enrichment of membrane proteins. (C) H_3_R and TfR expressions were quantified by densitometric analysis to assess differences in expression levels, and CCI animals showed a significant increase in H_3_R expression but not TfR expression compared with sham animals. The data are presented as the mean ± SEM. Statistical significance was assessed using unpaired Student’s t-test; **p < 0.01 N=5.

Overall, the expression results demonstrate that membrane-associated H₃R expression increases significantly in DRG tissue following peripheral nerve injury. Following the observation of increased H_3_R protein expression in the whole DRG tissue after nerve injury, we next sought to determine which DRG cell populations contribute to H3R levels by profiling Hrh3 expression at the single-cell level.

### 3.2 Assessment of H_3_R receptor expression in DRG at the single-cell level

Next, we examined the cell types expressing H_3_R to better understand its potential role in nociceptive signaling using publicly available GEO single-cell RNA-seq (scRNA-seq) datasets from mouse DRG. We analyzed the GSE174430 dataset, which provides comprehensive transcriptional profiles of all major DRG cell classes. Uniform manifold approximation and projection (UMAP) analysis suggested that DRG has six major cell types: neurons, satellite glial cells (SGCs), Schwann cells, fibroblasts, immune cells, and endothelial cells (**Figure 2A and 2B**). The expression analysis revealed that H_3_R expression was highly enriched in the DRG neuronal population compared to other cell types, with negligible expression in non-neuronal populations. Other histamine receptors (H_1_R and H_2_R) were detectable in neurons and immune cells (**Figure 2C and D**). As an additional comparison (data not shown), *Hrh3* was below the detection limit in the non-neuronal-cell dominant dataset GSE216665 from sciatic nerve. Furthermore, we also assessed expression of other histamine-related genes and neurotrophin-related genes involved in sensory neuron development and sensitization. Expression dot plot (**Figure 2D**) revealed that the Hdc gene, responsible for histamine synthesis, was highly expressed in immune cells, while the histamine metabolism enzyme gene, Maob, was only expressed in fibroblast and endothelial cells. Most neurotrophin receptors are expressed in neurons, fibroblasts, and SCG, except Ntrk1, which is expressed only in neuronal populations (**Figure 2D**).

**Figure 2.**
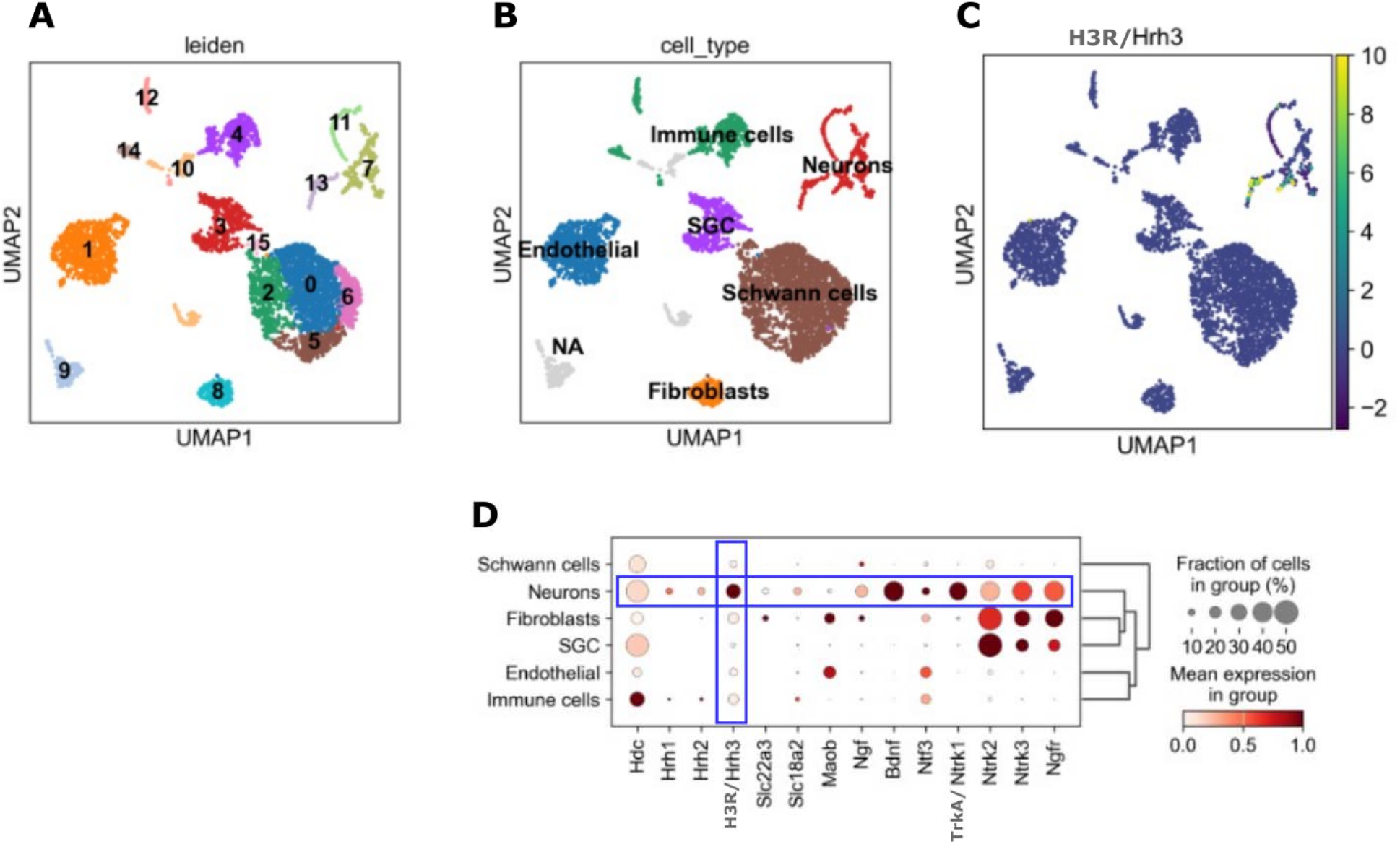
Histamine and neurotrophins-related gene expression across DRG cell types by transcriptomic profiling of the public GEO dataset. (A) UMAP showing 14 clusters based on a similar transcriptional profile, as determined by Leiden clustering. (B) A UMAP plot of different cell types of DRG, both neuronal and non-neuronal, with each cell type color-coded differently. (C) A UMAP plot featuring H_3_R transcripts across various DRG cell types showing enrichment of H_3_R in the neuronal population. (D) Dot plot represents the expression of various histamine and neurotrophin related genes across various annotated DRG cell types. Colors represent the mean expression scale. The sizes of the circles represent the percentage of cells in each specific cell type.

To further validate our RNAseq finding of H_3_R in neurons, we used cultured primary DRG neurons from neonatal mouse pups and performed immunofluorescence (IF) staining. The cellular identity of the cultured cells was confirmed prior to H_3_R staining using the anti-NeuN and anti-beta-3-tubulin antibodies as pan-neuronal markers, and anti-GFAP for satellite glial cells. IF staining with H_3_R antibody showed it to be expressed in mouse DRG culture neurons as indicated by co-localization of H_3_R with neuronal markers, including NeuN and beta 3 tubulin (tub-III) (**Figure 3A**). We did not observe any co-localization of H_3_R with glial cell marker GFAP in the DRG cell culture (**Figure 3A**). IF staining of adult rat DRG sections using NeuN and GFAP markers revealed a comparable expression pattern, with some level of H_3_R immunoreactivity also detected in glial cells (**Figure 3B**). As expected for a GPCR, H_3_R is primarily associated with the plasma membrane, although intracellular receptor pools may also be detected because of receptor trafficking, internalization, and recycling (Manchanda et al., 2024; Laniel et al., 2025). Some nuclear-associated staining was observed, but given the lack of established nuclear H_3_R localization, it may reflect nonspecific antibody reactivity. Overall, IF staining of both the DRG cultures and DRG sections is consistent with single-cell RNA-seq findings, supporting the presence of H_3_R expression primarily in neurons, with minimal to no expression in glial cells.

**Figure 3.**
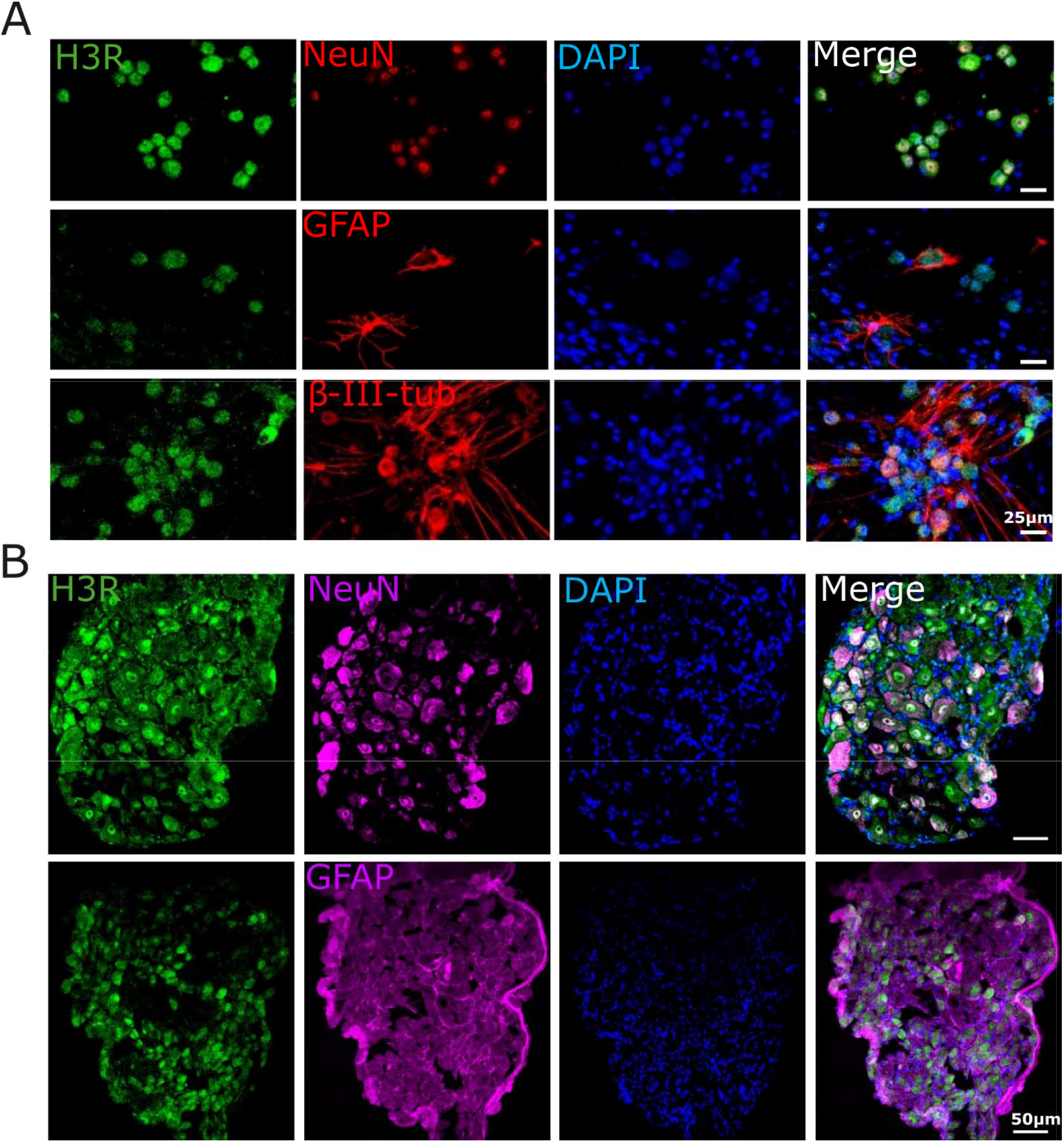
Localization of H_3_R expression in primary DRG cell culture and adult DRG tissue. (A) Representative immunofluorescent images showing H_3_R co-localization with neuronal marker (NeuN and βIII-tub) and the glial marker (GFAP) in isolated culture of mouse DRG cells. (B) Representative immunofluorescent images of rat DRG sections showing H_3_R co-labeled with NeuN and GFAP.

### 3.4 Transcriptomic profiling reveals DRG neuronal subtype expression of H_3_R

As our immunocytochemical findings and scRNA-seq analysis revealed, H₃R is specifically expressed in DRG sensory neurons, with very low/nearly undetectable levels in non-neuronal populations. We aimed to assess the sensory neuron subtype expressing H_3_R by reanalyzing publicly available scRNA-seq neuron specific datasets of mouse embryonic DRG (GSE139088) and adult DRG (GSE254789).

First, we investigated H_3_R expression in DRG sensory neuron subtypes in the GSE139088 embryonic dataset. This dataset contains well-annotated DRG neuronal subclasses, including CGRP, nonpeptidergic nociceptors, C-LTMRs, Aβ, and several other sensory neuronal subclasses. Expression UMAP analysis of H_3_R suggested that overall H_3_R expression was low and that a detectable H_3_R signal was not uniformly distributed across all DRG neuronal subtypes. Instead, H_3_R expression was preferentially associated with a limited subset of sensory neurons, most prominently CGRP⁺ peptidergic populations, including CGRP-α, CGRP-γ, and CGRP-ε neurons, as well as C-LTMR neurons but no other populations including nonpeptidergic, sst and other populations (**Figure 4 A, B**). Because peptidergic nociceptors are closely linked to NGF-TrkA signaling, a pathway strongly implicated in pain and peripheral sensitization, we next examined whether H_3_R-expressing neurons overlapped with NrtK1 sensory neurons at the single-cell level. Although H₃R and TrkA expression were detected in partially overlapping neuronal populations (**Figure 2B and C**), this did not establish whether they were expressed in the same cells. Therefore, we performed an overlap analysis using raw count data to identify cells with detectable expressions of both **H_3_R** and **Ntrk1**. Double-positive cells were defined as those with expression of both genes above zero counts and were visualized on the UMAP. Interestingly, we observed that subsets of H_3_R-expressing neurons overlapped with the TrkA+ neuronal population (**Figure 4D**). This was further confirmed through IF staining of cultured DRG neurons, which showed colocalization of H_3_R and TrkA (**Figure 4E**). These findings support the possibility that H_3_R contributes to subtype-specific modulation of peripheral sensory processing.

**Figure 4.**
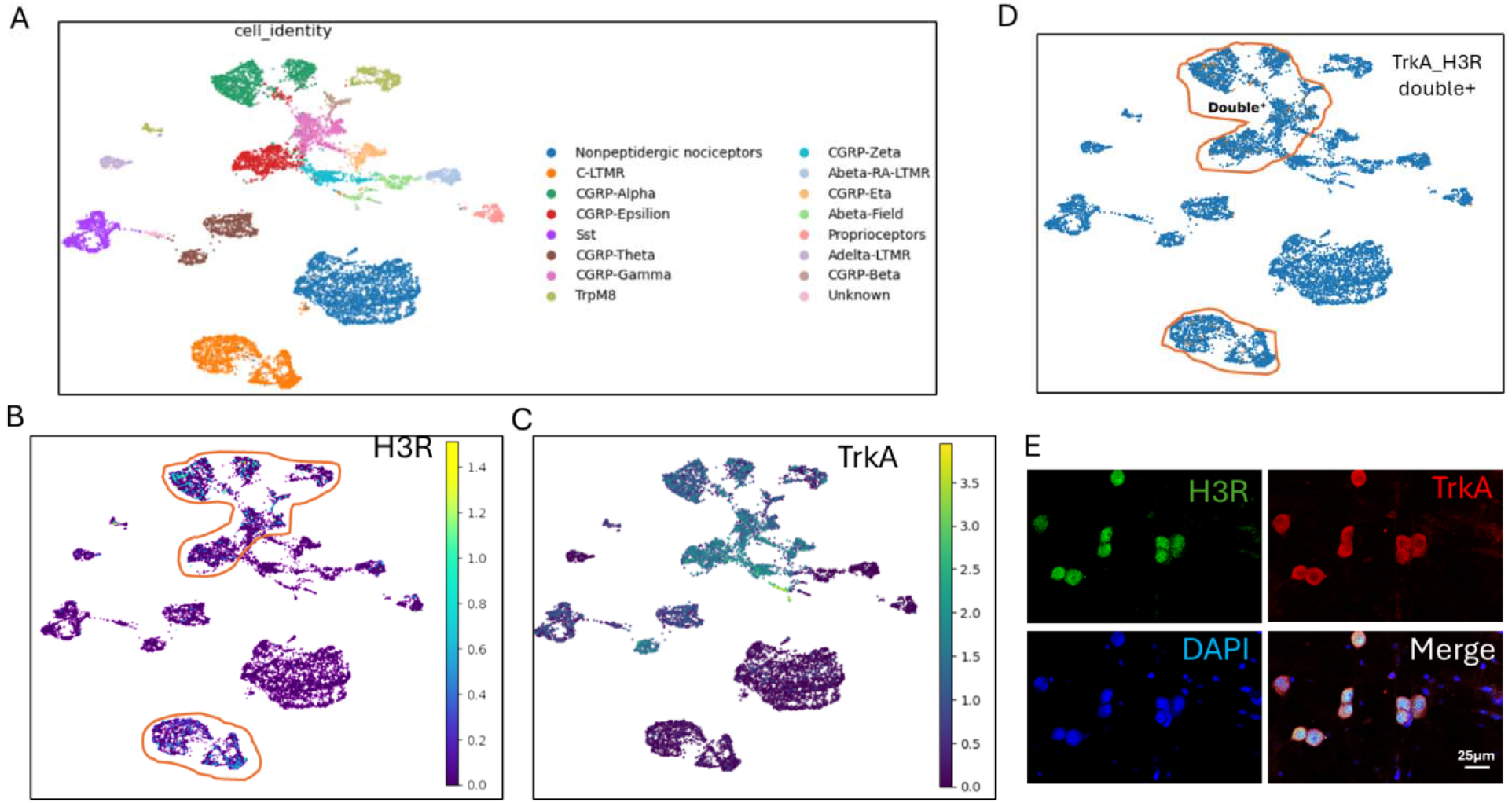
Transcriptomic profiling of H_3_R in embryonic DRG neurons subtypes and H3R IF imaging of cultured DRG neurons. (A) UMAP showing various annotated DRG neuronal subtype upon analysis of E11 DRG single-cell transcriptomic data from the GEO dataset (B) A UMAP plot featuring H_3_R transcription across DRG neuronal subtypes. (C) A UMAP plot featuring the NrtK1 transcripts distribution across DRG neuron populations. (D) UMAP showing a subset of DRG neurons with overlapping Nrtk1 and H_3_R molecular identities. (E) Representative immunofluorescence staining of DRG culture neurons having colocalized expression of H3R and TrkA. Scale bar:50 µm.

To extend our embryonic DRG neuron analysis to mature sensory neurons, we analyzed adult mouse sensory neurons in scRNA-seq datasets (GSE254789) and profiled H_3_R expression across defined mature sensory neuron subtypes, including peptidergic, non-peptidergic, C-LTMR, Aβ-LTMR, and pruriceptors. Analysis of the GSE254789 dataset showed that H_3_R expression was low but detectable only in neuronal populations. UMAP visualization revealed H_3_R-positive cells are primarily found within C-LTMR and peptidergic nociceptor populations, with sparse expression also observed in Aβ-LTMRs (**Figure 5A and B**). Interestingly, this distribution was consistent with our embryonic DRG neuron findings, which showed that H_3_R-positive sensory-neuronal subsets are associated with the TrkA-expressing lineage (**Figure 5A-C**).

**Figure 5.**
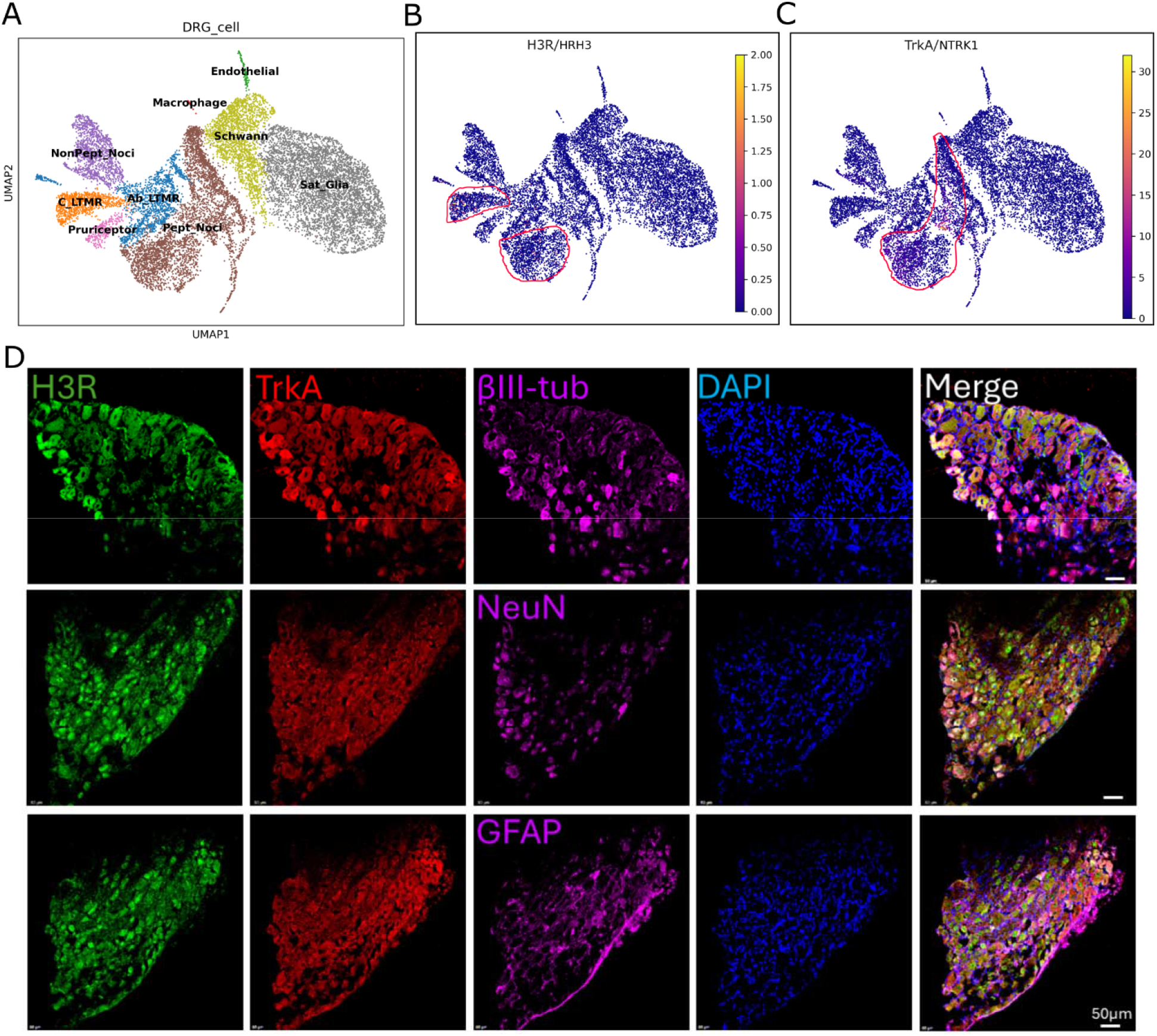
Expression of H_3_R and TrkA in neuronal and non-neuronal cells of adult DRG. (A) UMAP shows various annotated DRG neuronal and non-neuronal cells upon analysis of adult DRG single-cell transcriptomic data from the GEO dataset (B) A UMAP plot featuring H3R transcript distribution across DRG neuronal (indicated with red line) and non-neuronal cell types. (C) A UMAP plot featuring the NrtK1 transcripts distribution across DRG neuronal (indicated with red line) and non-neuronal populations. (D) Representative immunofluorescence images of adult rat DRG sections co-labeled for H_3_R and TrkA in neurons a marked by NeuN and tubb3, and GFAP as a satellite glial marker. Overlapping staining with NeuN indicates the presence of H_3_R in adult neurons with TrkA-positive sensory neuron identity. Scale bar: 50 µm.

We next used IF staining to assess H_3_R and TrkA expression in rat L4-L5 DRG sections to validate the neuronal specificity of H_3_R and its colocalization with TrkA in adult rat DRG tissue. To mark specific cell types in DRG, we used β-tub-III and NeuN as pan-neuronal markers and GFAP for satellite glial cells. The staining ascertained the neuronal identity of H_3_R by its colocalization with TrkA in β-tub-III and NeuN positive cells but very weak colocalization with GFAP, arguing against prominent H₃R expression in glial cells (**Figure 3B and 5D**). Additionally, we used other sensory neurons specific markers including CGRP as a marker of peptidergic nociceptive neurons, IB4 as a marker of small, unmyelinated C-fibers, and NF200 as a marker of large-diameter myelinated sensory neurons, including Aβ and Aδ fibers. IF staining of DRG sections revealed that H₃R and TrkA showed colocalization with the sensory neuron markers, with partial overlap with NF200⁺ neurons, stronger overlap in CGRP⁺ peptidergic neurons and variable but limited overlap with IB4⁺ neurons (**Figure 6**). This pattern is consistent with the sensory neuron specific transcriptomic distribution of H₃R in adult DRG, with the exception of variable overlap observed in IB4^+^ neurons (**Figure 5 A and B**).

**Figure 6.**
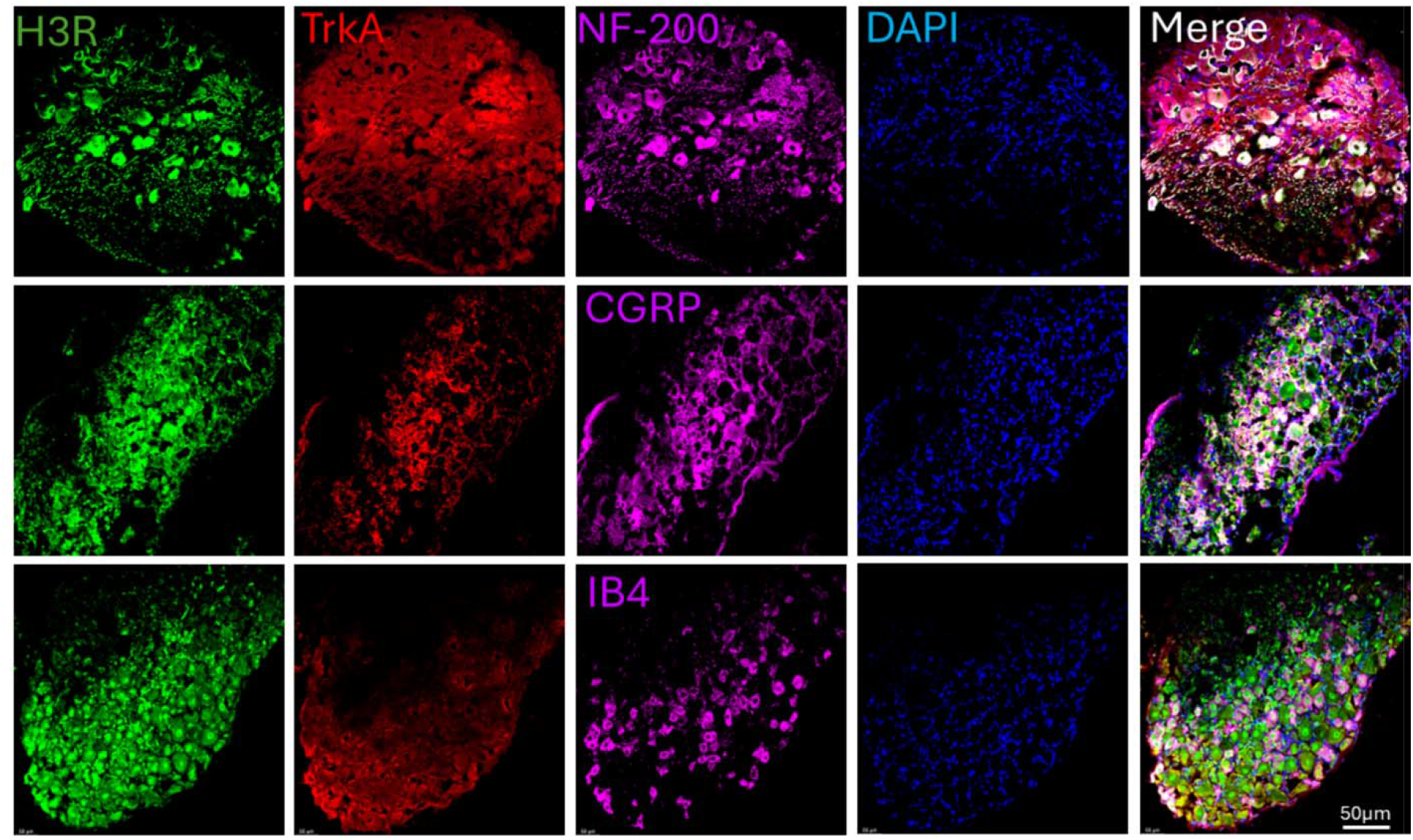
Immunostaining H_3_R and TrkA in rat DRG sections with sensory neuron markers. Representative immunofluorescent staining images showing expression of H_3_R and TrkA in sensory neuron subtypes through co-staining with NF-200, CGRP, and IB4 as markers of different sensory neuron subtypes. Scale bar:50 µm

Together, these findings establish that H₃R is broadly but selectively expressed across multiple DRG neuron subtypes with established roles in pain processing preferentially associated with sensory neuron populations involved in nociceptive sensitization, neuropeptide-mediated pain signaling, and mechanically evoked allodynia, while showing minimal association with glial populations.

### 3.5 Chronic antagonism of H₃R alleviates CCI-induced mechanical allodynia but not thermal pain

The upregulation of H_3_R expression in DRG tissues following CCI and its enrichment in C-LTMR, peptidergic, and a subset of Aβ DRG neurons suggest a potential role for H_3_R in the modulation of mechanical and heat-associated pain. We therefore sought to explore whether low doses of an H_3_R inverse agonist could alleviate neuropathic pain. Although several studies suggest that reduced nociception can be achieved through multiple mechanisms including regulating glial cell activation, our immunostaining and transcriptomic findings specifically showed H_3_R localization in neurons, with little or no detectable expression in glial cells.

To test this hypothesis, we used the treatment regimen described in **Figure 7A**. First, the acute effect of GSK administration (0.5 lllmg/kg, intraperitoneally) was assessed by comparing paw withdrawal thresholds in rats treated acutely with GSK compared to vehicle treated group following CCI. We found no difference in mechanical sensitivity between the GSK and vehicle treated groups, indicating that transient H_3_R inhibition is insufficient to alleviate neuropathic pain after CCI (**Figure 7B**, One-Way ANOVA; **p<0.01; N=6-7). Therefore, we administered GSK twice daily, starting on day 5 after CCI to assess its effect in neuropathic pain conditions. Mechanical thresholds were measured every second day, approximately 10-12 hours after GSK administration. Mechanical hypersensitivity reversed progressively, as indicated by a significant increase in paw withdrawal threshold values relative to CCI-vehicles from day 6 onwards, peaking at day 12 (**Figure 7C**, Two-Way ANOVA. **p < 0.01, ***p < 0.001, ****p< 0.0001 and #p<0.05 ##p < 0.01. N=12-13). This increase in threshold indicates that repeated drug administration is necessary for sustained anti-nociceptive effects. These data suggest that inhibition of H_3_R attenuates mechanical hypersensitivity in rats with neuropathic pain, underscoring the mechanosensory role of H_3_R.

**Figure 7.**
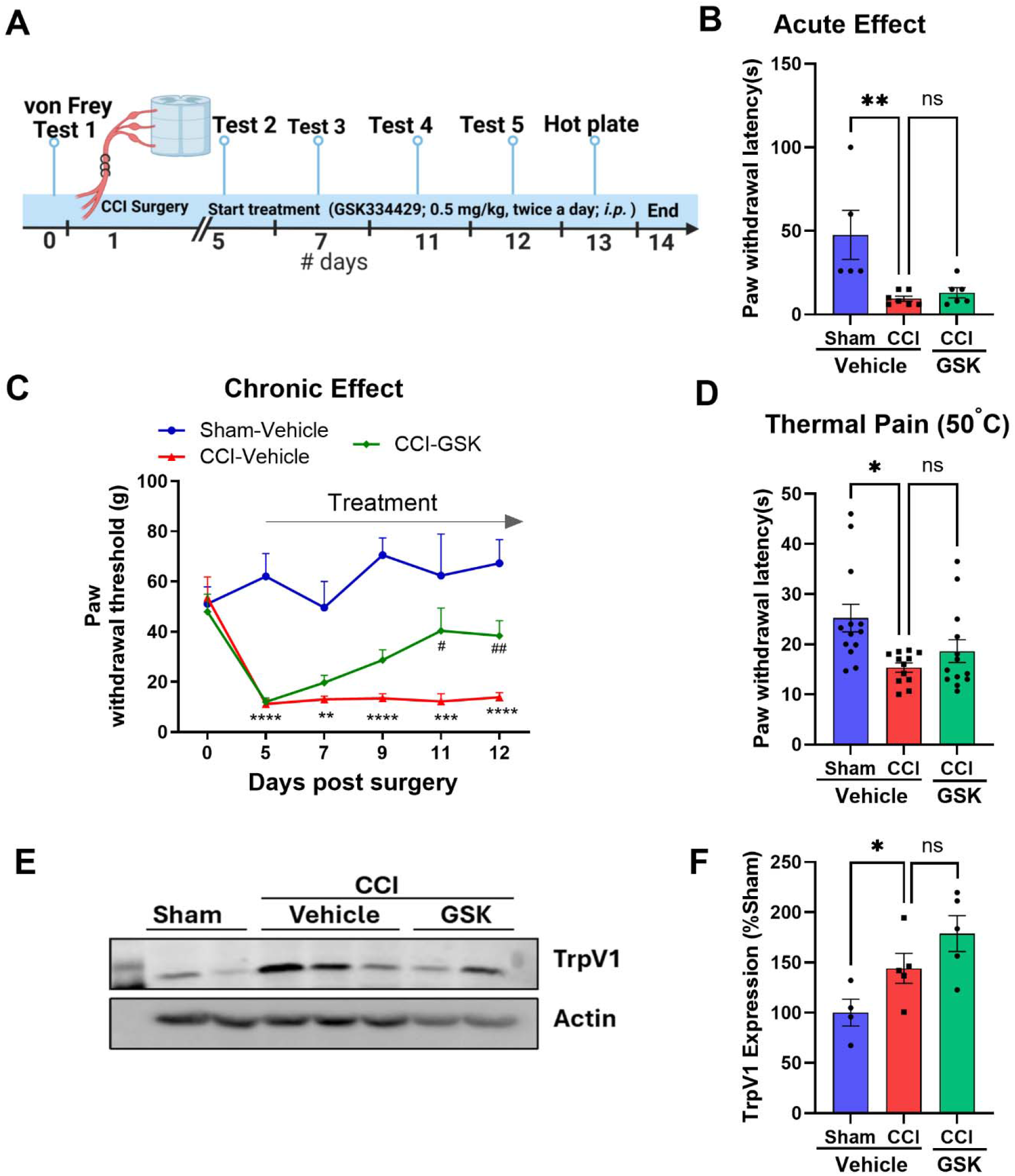
Treatment with H3R inverse-agonist alleviates CCI-induced mechanical pain but not the thermal pain. (A) *In vivo* experimental design and GSK treatment regimen (B) Acute effect of GSK (0.5 mg/kg) was evaluated 30 min after injection using the mechanical paw withdrawal threshold assay (One-Way ANOVA; \*\**p*<0.01; N=6-7). (C) Chronic treatment effect of H_3_R inverse agonist (GSK; 0.5 mg/kg twice a day), starting from day 5 after CCI, attenuated mechanical hypersensitivity compared with CCI-vehicle animals (Two-Way ANOVA; *F _(_*_2,164)_ =51.38, **p < 0.01, ***p < 0.001, ****p< 0.0001 and #p<0.05 and ##p < 0.01. N=12-13) (D) Animals treated with GSK (0.5 mg/kg), showed no alteration in the thermal pain sensitivity compared to CCI-vehicle controls, and both groups had significantly reduced paw threshold compared to the sham-vehicle controls as assessed by the hot plate test on day 12 at 50 °C. (Two-Way ANOVA. *p < 0.05, N=12-13) (E) Representative Western blots for TrpV1 expression in the spinal cord lysates from sham and CCI (vehicle and GSK) animals. β-actin represents a loading control. (F) TrpV1 expression was quantified by densitometric analysis to assess differences in expression levels, and CCI-vehicle animals showed a significant increase in TrpV1 expression compared with sham animals (One-Way ANOVA; \**p*<0.05; N=4-5). The data are presented as the mean ± SEM.

H_3_R antagonists have been shown to reduce thermal allodynia in the nerve injury model. Therefore, to further investigate the role of H_3_R signaling in thermal pain in our CCI animal model. We assessed thermal pain behavior and TRPV1 (a molecular sensor for heat) expression. The hot-plate test at 50 °C was used to evaluate thermal allodynia following CCI. There were no significant differences in the reaction times to thermal stimuli measured by paw licking or jumping between CCI animals administered the H_3_R inverse agonist and vehicle control groups (**Figure 7D**, One-Way ANOVA; **p<0.01; N=6-7).

To investigate the underlying molecular basis for this modality-specific effect, we examined the expression of transient receptor potential vanilloid 1 (TRPV1), a canonical heat-sensitive ion channel, in the lumbar spinal cord. Western blot analysis revealed that CCI significantly upregulated TRPV1 protein levels in the spinal cord compared to sham-operated rats (**Figure 7E & F**), confirming injury-induced thermal sensitization at the molecular level. However, chronic treatment with GSK did not alter the CCI-induced increase in TRPV1 expression, indicating that H₃R antagonism does not interfere with heat transduction mechanisms or thermal pain processing (**Figure 7E & F**, One-Way ANOVA; **p<0.01; N=6-7). These findings suggest that H₃R signaling selectively contributes to mechanical hypersensitivity, without affecting TRPV1-driven thermal pathways, and reinforce the therapeutic potential of H₃R inverse agonist/antagonists as mechanosensory-selective analgesics in neuropathic pain conditions.

### 3.6 Repeated H₃R antagonism reverses NGF-induced DRG neuron hyperexcitability

Transcriptomic analysis suggested that H_3_R is expressed in TrkA positive DRG neurons (**Figure 4 and 5**), implicating H_3_R signaling in NGF-responsive nociceptive populations. To evaluate the mechanistic role of H_3_R in NGF-driven modulation of neuronal hyperexcitability, we performed whole-cell patch-clamp (in current-clamp mode) recordings in primary DRG neurons following chronic NGF exposure with/without GSK as described in **Figure 8A**. Following treatment, the intrinsic excitability of DRG neurons was assessed by injecting a series of depolarizing current steps ranging from −50 pA to +140 pA in 10 pA increments. Long-term (48 hr) exposure of NGF (100ng/ml) significantly increases the frequency of evoked action potentials compared to vehicle treated controls across a broad range of current injections, specifically above 10 pA (**Figure 8 B and C**; Two-Way ANOVA, *p < 0.05, **p < 0.01, ***p < 0.001, ****p< 0.0001, N=15-25). This result confirmed that prolonged NGF exposure significantly enhances DRG neuronal excitability, consistent with previous reports of TrkA-mediated sensitization pathways (Mantyh et al., 2011; Donnelly et al., 2020; Moraes et al., 2022).

**Figure 8:**
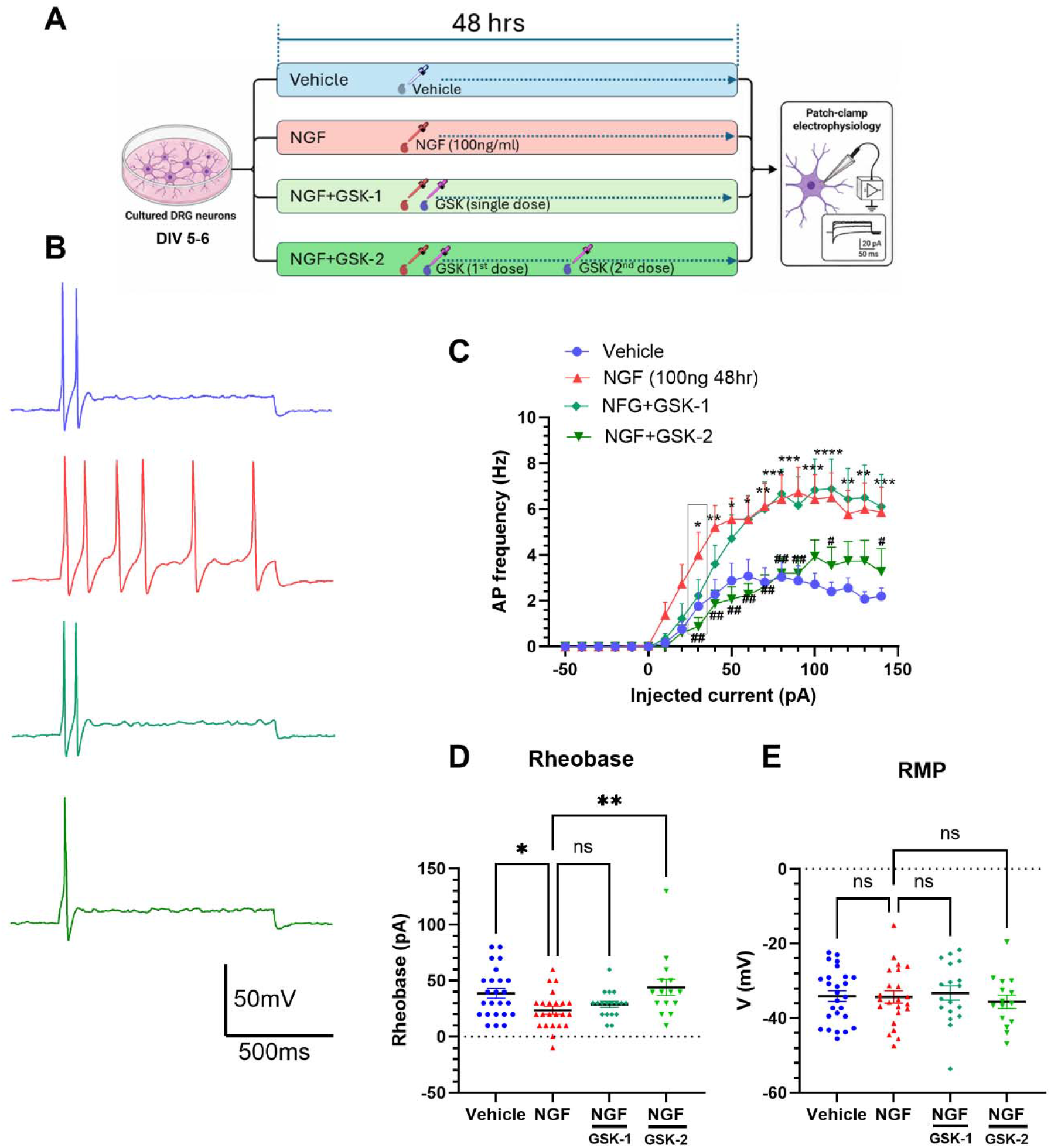
Sustained application of H₃R inverse agonist reverses NGF induced the hyperexcitability in primary DRG neurons. (A) Schematic of the experimental paradigm for *in vitro* treatment and patch-clamp electrophysiology. (B) Raw traces of evoked AP firing of cultured DRG neurons treated with vehicle (blue), NGF (red), NGF + GSK-1 (single dose of H₃R inverse agonist GSK334429,10 µM, day 1), and NGF + GSK-2 (two doses of GSK334429 day 1 and 2). Different colors represent different treatments of cultured DRG neurons. (C) Summary of evoked AP firing with repeated doses of H₃R inverse agonist reduced NGF-induced hyperexcitability reflected by AP firing of the DRG neurons (multiple comparison Two-Way ANOVA, * denotes Vehicle vs NGF; *p < 0.05, **p < 0.01, ***p < 0.001, ****p< 0.0001 and # denotes NGF vs GSK-2 #p<0.05, ##p < 0.01. N=15-25 neurons from three individual experiments). (D) Rheobase plot shows that less current is required in NGF-induced AP firing than in vehicle-treated neurons, which reverse with two doses of H_3_R inverse agonist (multiple comparison One-Way ANOVA, *p < 0.05, **p < 0.01, N=15-25 neurons from three individual experiments). (E) Resting membrane potential across various treatment conditions showed no significant changes between different groups. (One-Way ANOVA, p > 0.05, N=15-25). Data are presented as mean ± SEM.

To determine whether this hyperexcitation phenotype is modulated by H₃R activity, we applied GSK at a concentration of 10 μM under two dosing regimens: a single application at the start of NGF exposure (0 hr) and a double-application strategy, with the second dose administered at 24 hours (**Figure 8A**). Single application of GSK with NGF produced only a moderate and statistically nonsignificant effect on AP firing frequency, observed only at the lowest depolarizing step of 30 pA (**Figure 8C**). In contrast, repeated administration of GSK at 0 and 24 hours resulted in a significant suppression of NGF-induced hyperexcitability, effectively restoring AP firing frequency to levels comparable to control neurons across the 30–100 pA range (**Figure 8 C**; Two-Way ANOVA, #p<0.05, ##p < 0.01, N=15-25). This indicates that sustained H₃R inhibition is necessary to counteract the progressive changes in NGF induced excitability and may reflect the time-dependent nature of receptor engagement and downstream signaling cascades.

In addition to AP spike frequency, we also measured the rheobase, the minimum current required to elicit the first action potential. NGF exposed DRG neurons exhibited a significant reduction in rheobase compared to untreated controls and single dose of GSK application failed to restore rheobase values (**Figure 8D**; One-Way ANOVA, p > 0.05, N=15-25). As expected, the repeat application showed robust normalization of rheobase and significant change compared to the NGF alone, reaching levels statistically indistinguishable from those of control neurons (**Figure 8D**; One-Way ANOVA, *p < 0.05, **p < 0.01, N=15-25). Across all groups, none of the treatments significantly affected the resting membrane potential (RMP) of DRG neurons (**Figure 8 E**; One-Way ANOVA, p > 0.05, N=15-25), suggesting that observed changes in excitability occurred without a detectable alteration in basal membrane potential. Together, these findings demonstrate that NGF induces a hyperexcitable state in DRG neurons, characterized by increased firing frequency and decreased rheobase, and this sensitized state reversed by persistent H₃R blockade. These in vitro results complement our in vivo behavioral observations, where repeated but not acute H₃R inverse agonist treatment alleviated mechanical allodynia in CCI rats.

## 4. Discussion

The present study highlights a previously underappreciated role of H_3_R signaling in the pathophysiology of neuropathic pain following peripheral nerve injury. Our findings show upregulation of membrane associated H_3_R protein in neuropathic pain; negative modulation of the H_3_R receptor attenuates mechanical pain but not thermal pain. Transcriptomic data and immunofluorescence suggest H_3_R enrichment in TrkA^+^ peptidergic nociceptors and CLTMRs: cellular localization that links H_3_R to NGF-TrkA signaling and helps explain the modality selective (mechanical but not thermal) effects observed in behavior and electrophysiology. Overall, the results suggest a contributing rather than adaptive role for H_3_R in sensory neurons to elevated mechanical nociception following injury.

Multiple lines of evidence suggest that peripheral nerve injury alters the expression and membrane localization of multiple receptors and channels (Ji et al., 2002; Dai et al., 2004; Wood et al., 2004; Bray et al., 2013; Jang et al., 2017). Based on this, we investigated whether peripheral nerve injury affects H_3_R expression in the DRG. Like other membrane receptors that show altered expression during nerve injury, H_3_R membrane expression was upregulated after chronic constriction injury (Figure 1). Despite the implication of sensory H_3_Rs in nociception (Cannon et al., 2003, 2007; Hough and Rice, 2011), cell-type dependent expression of H_3_R in DRG remains poorly defined. To characterize cell specific expression of H_3_R, we re-analyzed the publicly available GEO scRNA-seq dataset GSE174430 from adult mouse DRG. We found that H_3_Rs are exclusively present only in the DRG neuronal population with minimal expression in other cell types, including glial cells, endothelial cells, and fibroblasts. In contrast some reports suggested that H_3_R is also present in glial cells (Iida et al., 2015; Xu et al., 2018; Degutis et al., 2025). Our immunohistochemical findings mostly support neuron-specific H_3_R expressions in DRG neurons. Immunofluorescence of early-stage DRG cultures and adult tissue sections confirmed the presence of H_3_R in sensory neurons with minimal expression in glial cells (Figure 3).

Embryonic (E11) and adult DRG neuron scRNA-seq datasets revealed H_3_R expressions mostly in peptidergic and C-LTMR classes of sensory neurons (Figure 4 and 5). Peptidergic neurons are mainly nociceptors marked by CGRP and often express TrkA receptors, whereas C-LTMRs are unmyelinated low-threshold mechanoreceptors. After nerve injury, C-LTMRs participate in mechanical pain. Both immunofluorescence and transcript-level findings in early stage and adult DRG tissue demonstrated that H_3_R co-expresses with TrkA at the level of individual neurons (Figure 3-5). TrkA is a high-affinity cell surface receptor of NGF and is crucial for the development and function of sensory neurons (Huang and Reichardt, 2003). After nerve injury, several inflammatory mediators including prostaglandin, interleukin, histamine, CGRP and NGF are released at injury sites and contribute to peripheral sensitization. NGF binds to the TrkA receptor and is retrogradely transported toward DRG. NGF-TrkA complex signaling increases sensitization by increasing ion channel expression in DRG sensory neurons (Chuang et al., 2001; Bonnington and McNaughton, 2003; Kitamura et al., 2005). Sensory neuron subtype-specific co-expression of H_3_R suggests a potential interaction between H_3_R and NGF pathways.

H₃R inverse agonist treatment reduced mechanical hypersensitivity in CCI animals without affecting thermal pain (**Figure 7**). Our scRNA-seq and immunohistochemical analyses showed that H₃R is enriched in CGRP positive peptidergic nociceptors and colocalizes with TrkA-positive DRG neurons. In contrast, H₃R expression was not detected in nonpeptidergic IB4⁺ or Sst⁺ neuronal populations in our dataset. Notably, the cross-species DRG transcriptomic atlas indicates that the CGRP-enriched TrkA^+^ peptidergic population has minimal expression of TRPV1 (Jung et al., 2023), which participate in the thermal nociceptive pathway. Consistent with this distinction, western blot analysis showed that TRPV1 remained upregulated following H₃R inverse agonist treatment, suggesting that TRPV1-associated thermal nociceptive pathways were not effectively modulated.

H_3_R exhibits constitutive activity and is coupled to G_i/o_-dependent pathways (Morisset et al., 2000); increased H_3_R expression could amplify downstream signaling mechanisms that promote neuronal hyperexcitability and pain hypersensitivity. Anti-nociceptive effects of H_3_R ligands have been pharmacologically demonstrated in multiple studies (Cannon et al., 2003; Hsieh et al., 2010; Łażewska and Kieć-Kononowicz, 2014; Mei et al., 2020; Obara et al., 2020; Stasiak et al., 2024). Electrophysiological recording demonstrated increased firing frequency and reduced rheobase currents in DRG cultured neurons exposed to NGF for 48 hours (Kitamura et al., 2005). This mimicked neuropathic pain-related hyperexcitability. In our study, the NGF-induced hyperexcitability was significantly dampened by sustained inverse agonism of H_3_R by selective ligand (GSK334429), with an insignificant effect from a single dose. This suggests a direct role for H₃R in regulating primary sensory neuronal excitability. The rescue of NGF-induced rheobase reduction and suppression of repetitive firing indicate stabilization of subthreshold membrane excitability at the nociceptor soma. Consistent with the possibility that injury or chronic NGF-exposure promotes a state where membrane-associated H₃R is increased, excessive receptor availability may shift H₃R signaling toward a maladaptive excitability-promoting state.

Although Gi/o activation typically suppresses excitability through direct activation of GIRK potassium channels and suppression of Ca^2+^ influx (Currie, 2010; Lüscher and Slesinger, 2010), some studies suggest excitatory effects of H_3_R activation in neurons (Rapanelli et al., 2016; Rivera-Ramírez et al., 2016; González-Sandoval et al., 2025). One plausible mechanism is that enhanced constitutive H₃R activity promotes G_i/o_ signaling leads to the steady dissociation of the heteromeric G-protein into G_i/o_ and free G_βγ_ subunit (Weis and Kobilka, 2018; Papasergi-Scott et al., 2024). G_βγ_-dependent activation of PLC_β_, promotes hydrolysis of membrane PIP₂, a critical cofactor for KCNQ channels and GIRK (Wettschureck and Offermanns, 2005; Suh and Hille, 2008; Falzone et al., 2026). H3R activity may cooperate with NGF-TrkA signaling which also activates PLC dependent hydrolysis of membrane PIP₂ (Chuang et al., 2001). Because PIP₂ is essential for KCNQ channel function, its depletion can suppress the M-current, thereby weakening an important constraint on neuronal excitability and promoting nociceptor sensitization (Zhang et al., 2003; Delmas and Brown, 2005; Linley et al., 2008). In this model, H₃R antagonist/inverse agonist treatment would limit excitatory H₃R signaling, preserve membrane PIP₂ availability, restore KCNQ channel function, increase rheobase, and suppress sustained repetitive firing. Thus, our data support injury-induced H₃R upregulation as a potential upstream contributor to NGF-driven DRG neuronal hyperexcitability. Ultimately, direct measurements of PIP₂ dynamics, KCNQ currents, or downstream Gi/o signaling will be needed to test this mechanism. Given the selective expression of H_3_R, cell-type specific manipulations, such as conditional knockouts or viral targeting, will be important for investigating the underlying mechanism.

Regardless of mechanism, our study identifies H₃R as a compelling therapeutic target for managing mechanical hypersensitivity in neuropathic conditions. The robust increase in membrane-localized receptor protein post-injury, coupled with its selective expression in NGF-responsive sensory neurons, provides a rational basis for selective pharmacological targeting. Moreover, our data suggest that modality specificity (mechanical vs. thermal) could be leveraged to avoid undesirable side effects, such as thermal hypoesthesia, often seen with broader analgesics. Future studies are required to determine the downstream signaling of H_3_R impacted by nerve injury.

In conclusion, the present study integrates transcriptomic, molecular, histological, behavioral, and electrophysiological approaches to demonstrate that H3R expression in sensory neurons is responsive to NGF and is upregulated in DRG neurons following peripheral nerve injury. Pharmacological inhibition of H3R reduces NGF- and injury-associated neuronal hyperexcitability and alleviates mechanical allodynia. Together, these findings identify neuronal H3R as a functionally relevant contributor to neuropathic pain and provide a foundation for developing cell-specific, targeted therapeutic strategies.

## Acknowledgements

This work was initiated at CSIR-CDRI Lucknow, India, where AK and SD received financial support from University Grants Commission (UGC) India. Subsequent experiments, data analysis, and preparation of the manuscript were completed at Washington University in St. Louis. We thank Prof. Valeria Cavalli for generously sharing GSK334429, NGF, anti-H_3_R and TrkA antibodies. We also thank Dr. Yukitoshi Izumi for generous support in this study.

## Contribution

AK contributed to conceptualization, data curation, visualization, and manuscript preparation. AK generated Figures 1, 2, 4, 5, 7, and 8; contributed to Figures 3 and 6; created the scientific illustrations; and wrote and edited the manuscript. RK contributed to figures 3, 4, 5, 6 and 8. PK contributed figure 1. AB contributed to figure 3, 5 and 6. AY contributed to figure 5, 6 and writing. BA, SD, and Deepmala contributed to Figure 1 and 8. All the authors participated in editing the manuscript. BC, CFZ, PNY and SM contributed to supervision, data interpretation, and manuscript review and editing.

## Declaration of generative AI and AI-assisted tool in the preparation of manuscript

During the preparation of this work the author(s) used ChatGPT and Microsoft Word grammar suggestions to ensure grammatical and stylistic flow. The author also used an AI tool to create graphical abstract. After using these tools, the authors reviewed and edited the content as needed and take full responsibility for the content of the publication.

