## Supplemental file for "Targeting Histamine H₃ Receptors to Suppress NGF-Driven Hyperexcitability in Neuropathic Pain"

Figures: Images of raw western blot

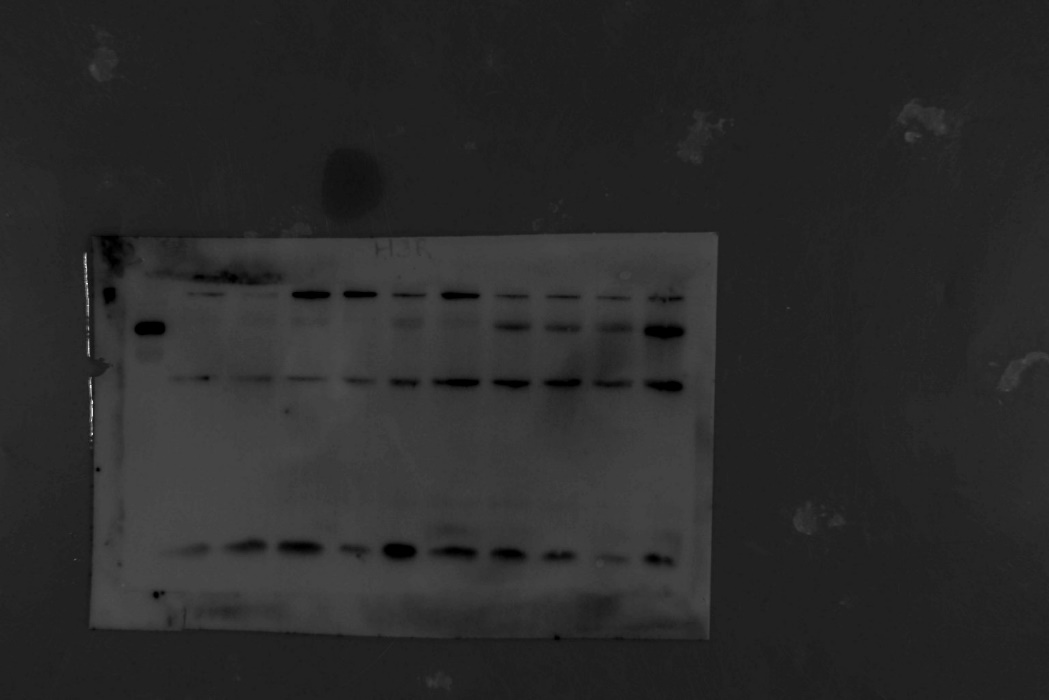

Figure1:H3R WGA blot

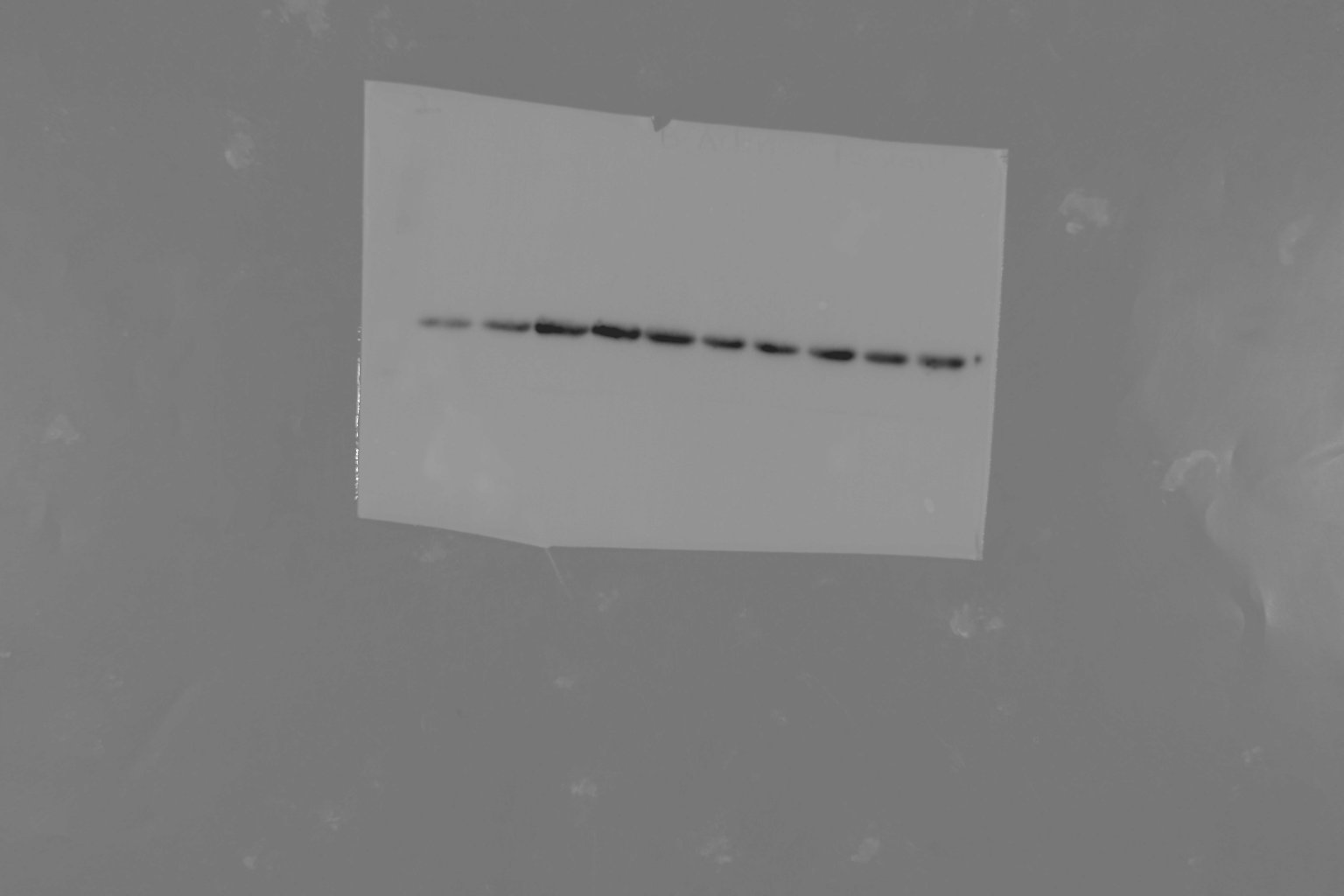

Figure1: Actin blot

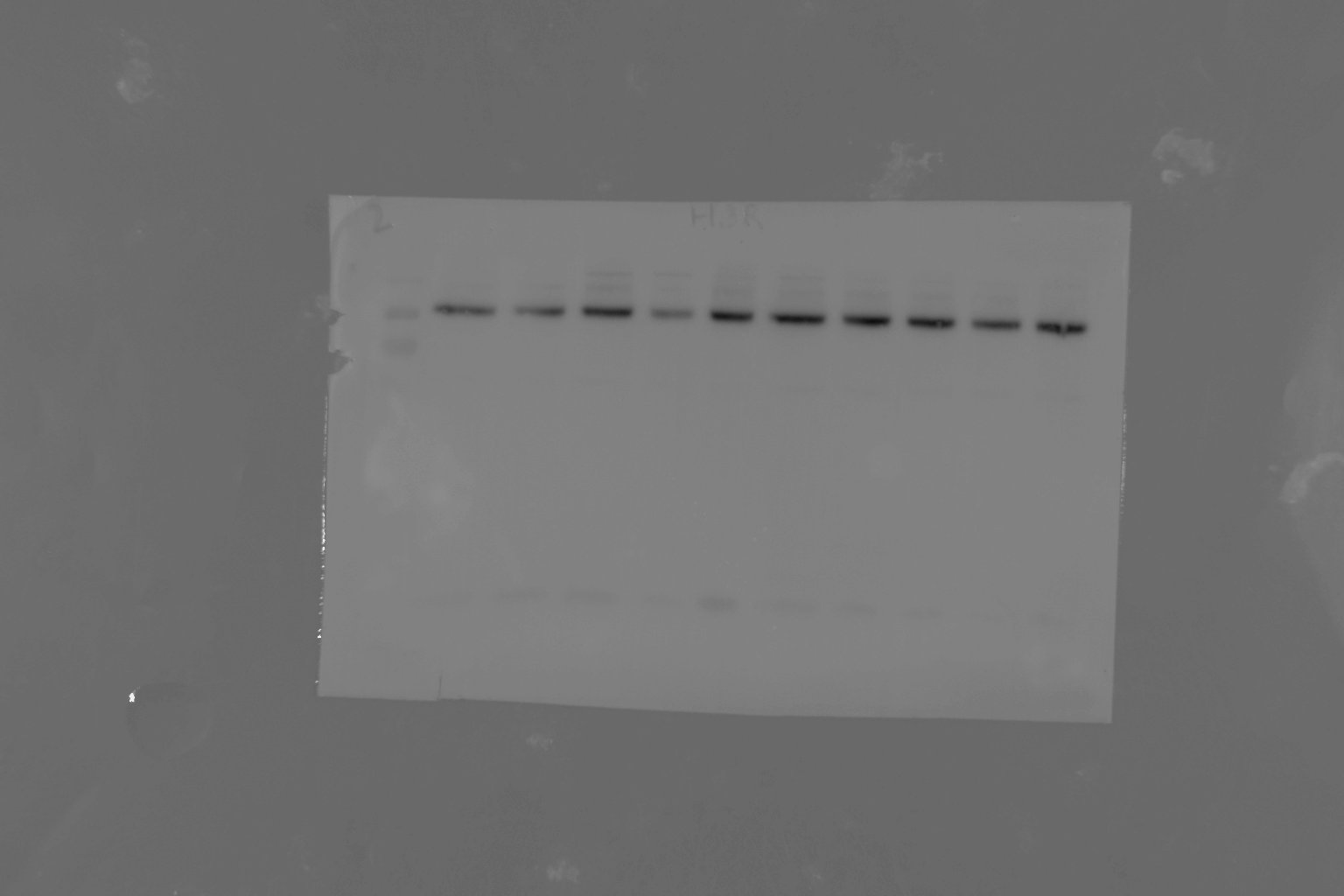

Figure1: Transferrin WGA blot

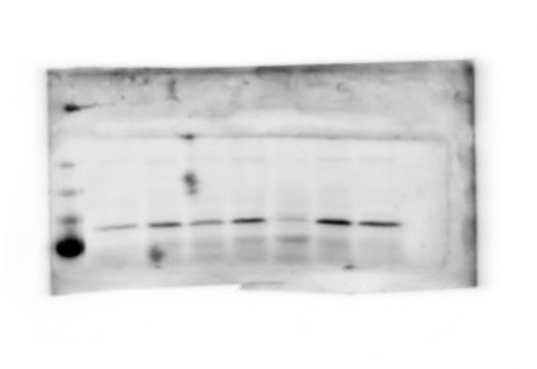

Figure7F: TrpV1 blot 1, used in quantification

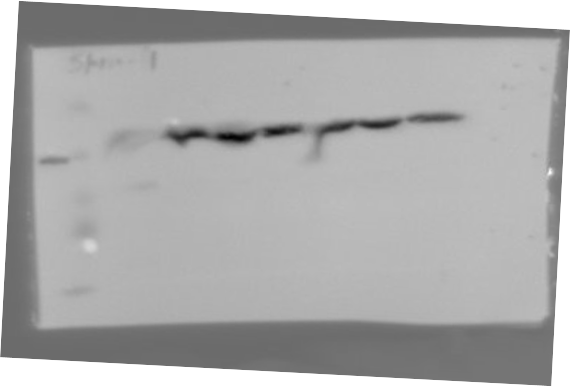

Figure 7F: Actin blot 1, used in quantification

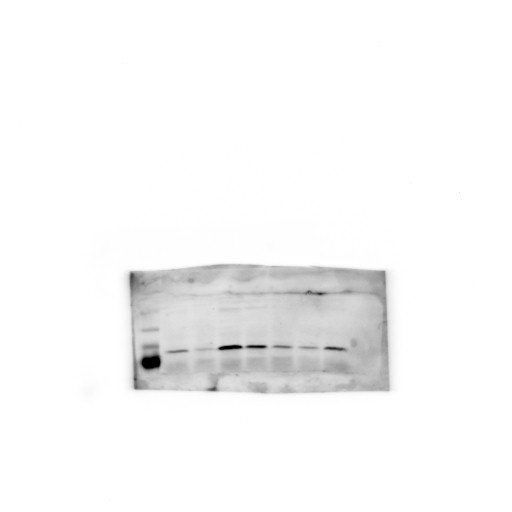

Figure7E: TrpV1 blot 2, used in quantification

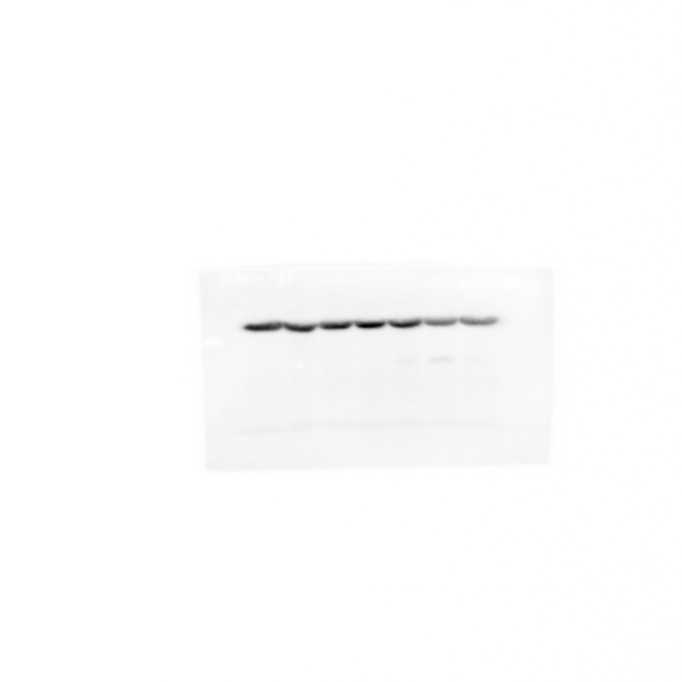

Figure 7E: Actin blot 2, used in quantification
